# The role of m6A RNA methylation in the maintenance of X-chromosome inactivation and X to autosome dosage compensation in early embryonic lineages

**DOI:** 10.1101/2024.04.22.590572

**Authors:** Hemant C Naik, Runumi Baro, Amritesh Sarkar, M Muralidhar Nayak, Kartik Sunagar, Srimonta Gayen

**Author notes:** Equal contribution (Hemant C Naik, Runumi Baro).

## Abstract

In therian mammals, inactivation of one of the X-chromosomes in females balances the dosage of X-linked gene expression between the sexes. On the other hand, upregulation of active-X balances the dosage of monoallelically expressed X-linked genes with biallelic autosomal genes (AA). Factors and mechanisms involved in the maintenance of X-chromosome inactivation (XCI) and X to autosome dosage compensation remain underexplored. Recently, it has been implicated that N6-methyladenosine (m6A) RNA modification contributes to XCI and X to autosome dosage compensation. Here, we have investigated the role of m6A RNA methylation in the maintenance of XCI and X to autosome dosage compensation in early embryonic lineages. Surprisingly, we find that the depletion of m6A RNA methylation does not affect the maintenance of inactive-X gene silencing in mouse epiblast stem cells (EpiSC), trophoblast stem cells (TSC) and extraembryonic endoderm stem cells (XEN). On the other hand, we show that m6A marks are less enriched on X-linked transcripts than the autosomal transcripts in early embryonic lineages. It is believed that less enrichment of m6A in X-linked transcript increases the stability of the X-linked transcript and thereby contributes to the X to autosome dosage compensation. Interestingly, we find that while X-linked transcripts without m6A are fully dosage compensated, transcripts with m6A undergo partial X to autosome dosage compensation in EpiSC, TSC and XEN. However, we find that the depletion of m6A has a minor effect on the X to autosome dosage compensation. Taken together, our study provides significant insight into the role of m6A RNA methylation in dosage compensation of early embryonic lineages.

## Introduction

In the course of evolution, the Y-chromosome underwent massive degradation through gene-loss events (Bachtrog, 2013; Graves, 2016; Lahn & Page, 1999). This rendered male-specific monoallelic expression of X-linked genes in contrast to biallelic expression from the autosomes. Into this, Susumu Ohno hypothesised that X-chromosome in males underwent upregulation to balance the X-to-autosome dosage (Ohno, 1967). Subsequently, the inheritance of this upregulated-X in females is thought to have created an overdose of X-linked gene products, which was rectified by the evolution of X-chromosome inactivation (XCI) mechanisms (Lyon, 1961). In recent years, the mechanisms of XCI have been investigated extensively. However, the mere existence of X-chromosome upregulation (XCU) in mammals has been contested for decades. Some studies refuted Ohno’s hypothesis (Chen et al., 2020; F. Lin et al., 2012; M. Wang et al., 2017; Xiong et al., 2010; Yang & Chen, 2019). On the other hand, many studies provided evidence in support of XCU (Borensztein et al., 2017; Cidral et al., 2021; De Mello et al., 2017; X. Deng et al., 2011, 2013; Di & Disteche, 2006; Gupta et al., 2006; Kharchenko et al., 2011; Larsson et al., 2019; H. Lin et al., 2007, 2011; Mahadevaiah et al., 2020; Mandal et al., 2020; Sangrithi et al., 2017; F. Wang et al., 2016; Yildirim et al., 2012). Importantly, recent studies leveraging allele-resolved transcriptomic analysis at the single-cell level have clearly shown coordinated upregulation of the active X-chromosome upon silencing of genes on the inactive X during early embryogenesis (Lentini et al., 2022; Naik et al., 2022, 2024). Together, while XCI balance the dosage of X-linked gene expression between sexes, XCU contributes to balancing the X to autosome dosage. However, the mechanisms and factors involved in the maintenance of XCI and X to autosome dosage compensation remain poorly understood.

N6-methyladenosine (m6A) is one of the most prevalent internal reversible modification known to be present on RNAs (Dominissini et al., 2012; Meyer et al., 2012). Recently, m6A RNA-methylation has emerged as a new player in gene regulation, which plays key roles in embryonic development, cellular differentiation, maintenance of cellular integrity and stress response (Batista et al., 2014; Geula et al., 2015; H. Lee et al., 2019; H. B. Li et al., 2017). Dysregulation of m6A dynamics has been attributed to many pathophysiological conditions, including cancer (Berulava et al., 2020; Jiang et al., 2021; Pupak et al., 2022). m6A modification is deposited on the RNAs by the core methyltransferase complex consisting of METTL3, which is the main catalytic methylase (writer), METTL14 and WTAP, along with accessory proteins (Liu et al., 2013; Ping et al., 2014). On the other hand, FTO and ALKBH5 act as an erasure for m6A RNA modification (Jia et al., 2011; Zheng et al., 2013). Finally, the ‘reader’ enzymes recognize the m6A mark on the RNAs and decide their fate within a cell (Knuckles et al., 2017; Shi et al., 2017). Several studies have shown that m6A deposition occurs mostly at the 5’- and 3’-regions of transcripts and is believed to regulate translation efficiency, alternative splicing events, nuclear mRNA export and mRNA turnover (Ke et al., 2015; Roundtree et al., 2017; Slobodin et al., 2017; X. Wang et al., 2013, 2015; Xiao et al., 2016). Although m6A-mediated regulation is known to majorly control RNA metabolism, recently, it has also been shown to act as a global epigenetic regulator (Y. Li et al., 2020; Selmi & Lanzuolo, 2022; Wei et al., 2022). Together, m6A RNA methylation has emerged as one of the key players in epigenetic regulation of gene expression. Emerging studies indicate that m6A could be a critical player in regulating dosage compensation as well. However, the precise role of m6A RNA methylation in dosage compensation remains underexplored. Into that, X-inactive specific transcript or the *XIST* lncRNA, which is the master regulator of XCI, has been shown to be highly enriched with m6A modification. Although some studies have implicated the role of m6A-regulation in *XIST*-mediated gene silencing (Chang et al., 2022; Nesterova et al., 2019; Patil et al., 2016), the essentiality of m6A in the maintenance of X-chromosome inactivation remains unknown. On the other hand, upregulation of the active-X has been reported at the transcriptional level through enrichment of active chromatin marks, increased chromatin accessibility and increased transcriptional burst kinetics of X-linked transcripts (X. Deng et al., 2013; Larsson et al., 2019; Talon et al., 2021). Given the versatile nature of m6A acting as a global epitranscriptomic regulator, it is thought that m6A-mediated regulation of transcript stabilisation contributes to equilibrate the X to autosome dosage. Indeed, a recent study has shown that m6A RNA-methylation contributes to the X to autosome dosage compensation (Ruckle et al. 2023). However, the potential contribution of m6A in fine-tuning X to autosome dosage across different cell types and developmental contexts remains underexplored. In this study, we have explored the relative contribution of m6A in the maintenance of XCI as well as in maintaining an equilibrium X to autosome dosage in different early embryonic lineages.

## Results

### Depletion of m6A RNA methylation does not perturb the maintenance of X-linked gene silencing in XEN, TSC and EpiSC

Mouse undergo two waves of XCI – imprinted inactivation (iXCI) of the paternal-X at the 2-4 cell stage of embryogenesis, followed by subsequent reactivation of the paternal-X in the inner cell mass cells of the late blastocyst and finally random inactivation (rXCI) of either of the paternal- or maternal-X in the embryonic epiblast (Harris et al., 2019; Huynh & Lee, 2003; Maclary et al., 2014; Okamoto et al., 2004; Takagi & Sasaki, 1975). To investigate the role of m6A RNA methylation in the maintenance of XCI, we used stem cell lines representing early embryonic lineages. We have used extra-embryonic endoderm (XEN) cells that represent the primitive endoderm of late blastocysts, which have already initiated their iXCI process and, thereby, maintaining the iXCI state (Fig. 1A). On the other hand, we have used trophoblast stem cells (TSC) that represent the trophectodermal cells of the blastocyst and are in a maintenance phase of iXCI (Fig. 1A). Apart from these two, we have used epiblast stem cells (EpiSC) lines resembling much of post-implantation epiblast, which represents the maintenance phase of rXCI (Fig. 1A). Notably, all are hybrid cell lines carrying polymorphic X-chromosomes (X^Mus^: *Mus musculus* origin and X^Mol^: *Mus molossinus* origin) and in all cell lines, the X^Mus^ is selectively inactivated, which allowed us to disentangle the X-linked gene expression between active vs. inactive X through allele-specific analysis (Fig. 1A). Previously, it was implicated that m6A RNA modification in Xist RNA help in Xist mediated gene silencing (Chang et al., 2022; Patil et al., 2016). Therefore, we delineated if m6A RNA modification is present in the Xist RNA expressing in XEN, EpiSC and TSC through performing methylated RNA immunoprecipitation sequencing (MeRIP-seq). We find that the m6A modifications are present at exon1 and exon7 of Xist RNA in TSC, XEN and EpiSC (Fig. 1B). Next, we depleted m6A in XEN, TSC and EpiSC through inhibition of m6A methyl transferase METTL3 using STM2457 inhibitor (Yankova et al., 2021) (Fig. 1C). To validate the m6A depletion upon inhibition of METTL3, we performed liquid chromatography-tandem mass spectrometry (LC-MS/MS) to quantitate m6A levels in mRNAs isolated from DMSO treated and inhibitor (STM2457) treated XEN, TSC and EpiSC cells. We observed robust m6A depletion upon inhibition of METTL3 (XEN, n=3; EpiSC and TSC, n=2) (Fig. 1C). Notably, in the inhibitor-treated cells, we observed a 76.5% reduction of m6A/A levels in XEN, 75.5% in TSC and 70% in EpiSC cells compared to DMSO treatment (Fig. 1C). Next, we performed RNA sequencing (RNA-seq) of DMSO vs. STM2457-treated XEN, TSC and EpiSC cells. From the RNA-seq data, we compared the Xist expression level between DMSO vs. inhibitor-treated cells. We find that Xist RNA exclusively express from the X^Mus^ allele in all three cell types, thereby validating that X^Mus^ is the inactive-X (Fig. 1D). We find that overall Xist expression remains unaltered between DMSO vs. inhibitor-treated TSC and EpiSC (Fig. 1D and 1E). However, we observed that Xist expression was reduced in XEN cells upon inhibitor treatment, although it was not statistically significant (Fig. 1D and 1E). Together, we conclude that m6A depletion does not affect the Xist expression level in TSC and EpiSC, however, it leads to the reduction in Xist expression in XEN cells. Next, we explored if m6A depletion affect the X-linked gene silencing. To test this, we performed allele-specific expression analysis of the RNA-seq data. We observed exclusive expression of X-linked genes from the active-X (X^Mol^) in all XEN, TSC and EpiSC (Fig. 1F). Upon inhibitor treatment, there were no overall changes in inactive-X expression compared to the DMSO-treated cells, indicating no effect on X-linked gene silencing despite the m6A depletion (Fig. 1F). To probe this further, we performed a chromosome-wide expression analysis of all the X-linked genes in XEN, TSC and EpiSC at an allelic resolution (Fig. 2A). As expected, we observed majority of the X-linked genes present on the inactive-X (X^Mus^) remained silenced in the DMSO treated XEN, TSC and EpiSC (Fig. 2B). Only a few genes showed constitutive biallelic expression which are known escapees of XCI (Berletch et al., 2011) (Fig. 2B). Surprisingly, we observed absolutely no signs of reactivation of inactivated genes upon m6A depletion through our allele-specific RNA-seq analysis in the inhibitor-treated XEN, TSC and EpiSC cells (Fig. 2B). Only, in TSC we observed reactivation of few genes (Ube2a, Nkap, Xlr3C, Usp11, Mmgt1, 8030474K03Rik). In contrast, we also observed inactivation of previously active genes in TSC (Praf2) and XEN (Rbm41). Overall, our analysis suggests that m6A RNA methylation is dispensable for maintaining the X-linked gene silencing in female XEN, TSC and EpiSC cells (Fig. 2C).

**Fig. 1:**
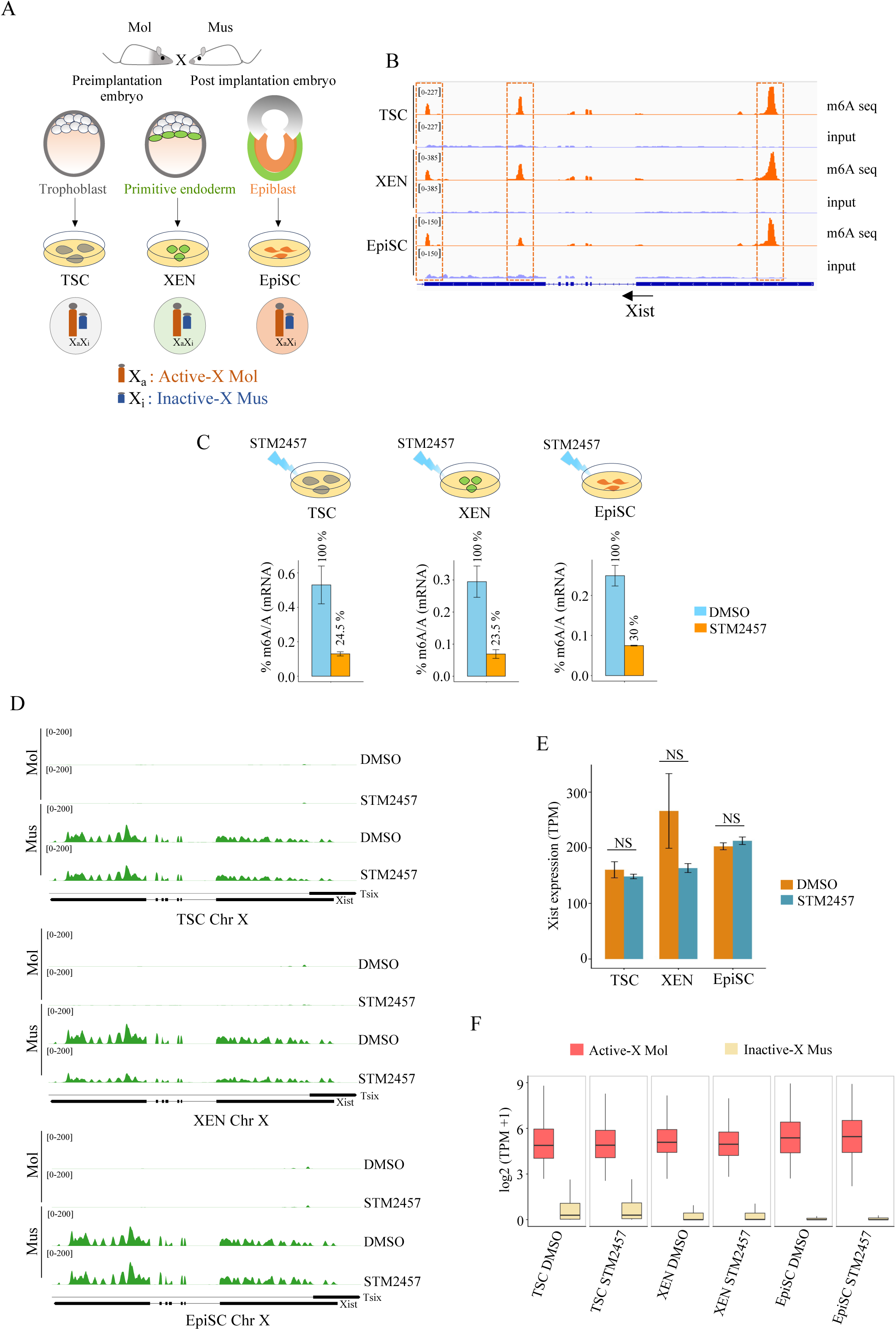
Profiling the effect of m6A depletion on Xist expression and maintenance of X-linked gene silencing in female TSC, XEN, and EpiSC. (A) Schematic showing the hybrid TSC, XEN, and EpiSC lines carrying polymorphic X-chromosomes (X^Mus^: *Mus musculus* origin and X^Mol^: *Mus molossinus* origin). X^Mus^ is inactivated in these cells. (B) IGV browser snapshot of m6A sites in Xist in TSC, XEN and EpiSC; dotted lines represent m6A sites. (C) Quantification of m6A level relative to A in mRNA of TSC, XEN and EpiSC by liquid chromatography-tandem mass spectrometry (LC-MS/MS) upon METTL3 inhibitor (STM2457, 50μM) and vehicle (DMSO) treatment for 12 hrs. Error bar represents standard deviation of mean for TSC (n=2), XEN (n=3), and EpiSC (n=2). (D) Tracks showing the allelic Xist expression in DMSO and STM2457 treated TSC, XEN, and EpiSC. (E) Quantification of Xist expression in DMSO vs. STM2457 treated TSC, XEN, and EpiSC; Two sample t-test, NS-non-significant. (F) Boxplot showing the expression of X-linked genes from active-X (X^Mol^) and inactive-X (X^Mus^) in DMSO vs. STM2457 treated TSC, XEN, and EpiSC. XEN – extraembryonic endoderm cells, TSC – trophoblast stem cells, EpiSC – epiblast stem cells.

**Fig. 2:**
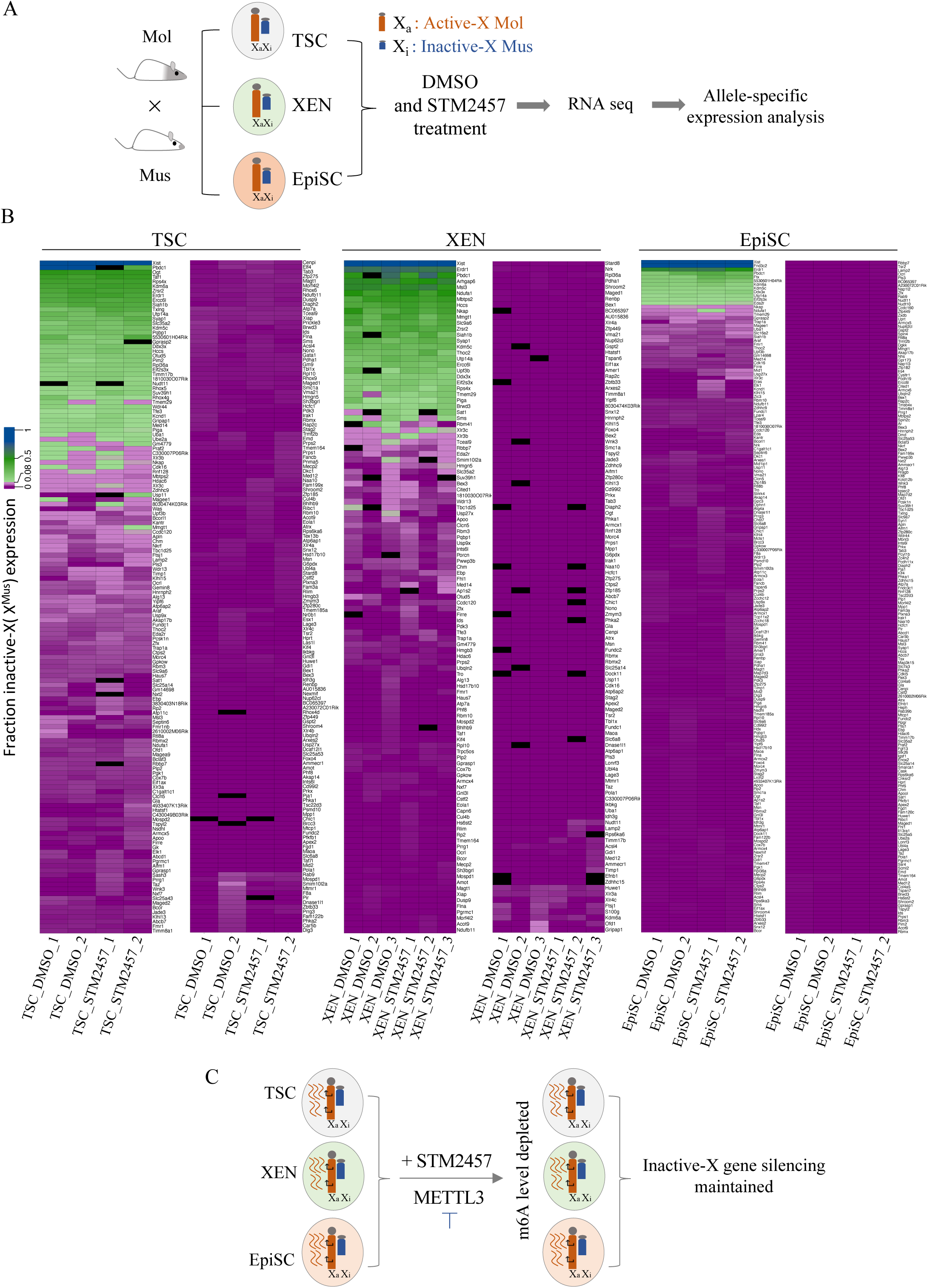
Depletion of RNA m6A level does not affect the maintenance of XCI. (A) Schematic workflow showing RNA-seq analysis in the DMSO and STM2457 treated TSC, XEN, and EpiSC at allelic resolution. (B) Heat map showing the fraction of expression of X-linked genes from inactive-X (X^Mus^) in DMSO vs. STM2457 treated TSC, XEN, and EpiSC. (C) Model diagram showing that despite of m6A depletion X-linked gene silencing on the inactive-X is maintained in STM2457 treated TSC, XEN and EpiSC.

### m6A is less enriched in X-linked transcripts compared to the autosomal transcripts in early embryonic lineages

Differential m6A enrichments in the transcripts is known to modulate the transcripts stability and balance the gene expression stoichiometry (Zaccara & Jaffrey, 2020). One of the prevailing hypotheses is that the higher stability of X-chromosomal transcripts relative to autosomal transcripts partially contributes to the balance of the X to autosome expression dosage in cells. Recently, it has been shown that less enrichment of m6A in X-linked transcripts than the autosomal transcripts renders to the better stability of X-linked transcript (Rücklé et al., 2023). However, the pattern of m6A enrichment in X-linked and autosomal transcripts during early embryogenesis remains unknown. To explore this, we profiled the m6A enrichment on X-linked vs. autosomal transcripts during early embryogenesis using available MeRIP-seq datasets (Wang et al., 2023) (Fig. 3A). Interestingly, we find that in GV oocyte m6A modifications are less enriched in X-linked transcripts compared to the autosomal transcripts (Fig. 3A). However, we did not find any significant differences in X-linked vs. autosomal m6A enrichments in MII-oocyte and Zygote (Fig. 3A). But, we observed that enrichment of m6A in X-linked transcripts is reduced from 2-cell stage to 8-cell and blastocyst stage compared to the autosomal transcripts (Fig 3A). Next, we extended our analysis to different embryonic lineages. For this, we performed MeRIP-seq in TSC, XEN and EpiSC lines and found that m6A modification is significantly less enriched in X-linked transcripts compared to the autosomal transcripts in these cells as well (Fig. 3B). Notably, we found that the enrichment of m6A level in the X-chromosomal transcripts are lesser than autosomal transcripts in different mouse embryonic stem cells (ESCs) lines and in embryoid body (Fig. 3C). Next, we extended our analysis to different somatic cell types. We observed the same trend for mouse embryonic fibroblast (MEF), neural progenitor cells (NPC), immature dendritic cells (imDC), mature dendritic cells (maDC), regulatory dendritic cells (regDC), and mesenchymal stem cells (MSC) (Fig. 3D). In contrast, of the three tissue types that we analyzed, difference in m6A levels was not observed in midbrain and hippocampus (6 weeks), however, m6A was lower on X-linked transcript compared to the autosomes in hippocampus (2 weeks) and liver tissue (Fig. 3D). Taken together, we conclude that m6A are less enriched in X-linked transcripts compared to the autosomal transcripts in early embryonic lineages as well as in somatic lineages, with exceptions to certain somatic tissues.

**Fig. 3:**
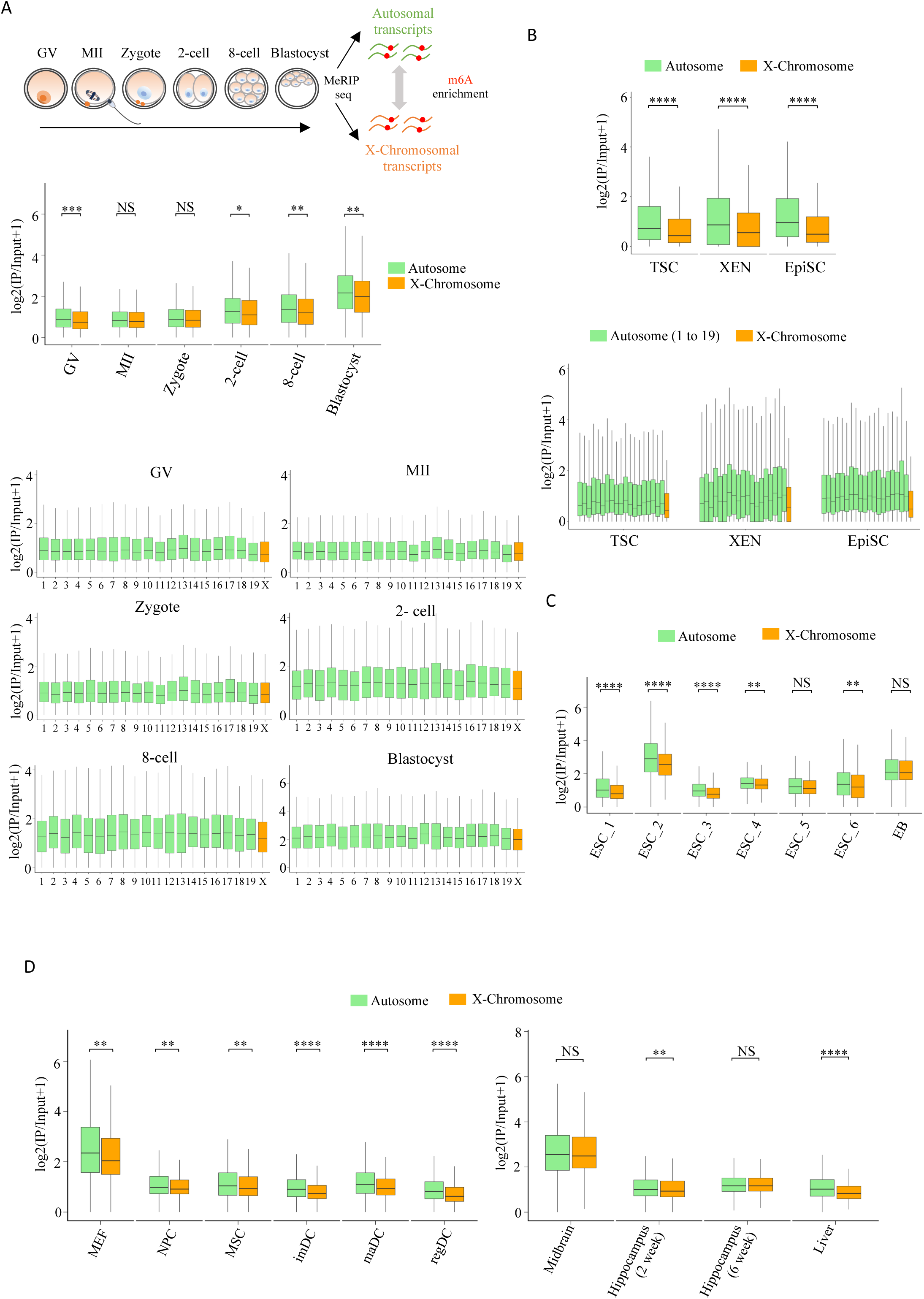
Profiling m6A enrichment between autosomal and X-chromosomal transcripts at different stages of early embryogenesis and different embryonic lineages. (A) Schematic showing the workflow of m6A enrichment analysis in autosomal and X-linked transcripts during early embryogenesis (top); RNA m6A enrichment autosomes vs. X-linked transcripts in mouse oocyte and various stages of mouse early embryonic development. Edges of each box represent the 25th and 75th quartiles, and the center line denotes the median value (middle); Mann-Whitney U test; *P value < 0.05, ** P value < 0.01, ***P value < 0.001, NS-nonsignificant; RNA m6A enrichment chromosome wise (bottom); GV-germinal vesicle stage oocyte, MII-metaphase II stage oocyte; (B) RNA m6A enrichment in autosomal vs. X-chromosomal transcripts in TSC, XEN, and EpiSC (top). Mann-Whitney U test, ****P value < 0.0001; Chromosome wise RNA m6A enrichment in TSC, XEN, and EpiSC (bottom). (C) Autosome vs. X-chromosomal transcripts RNA m6A enrichment in mouse ESC and EB; Mann-Whitney U test, **P value < 0.01, ****P value < 0.0001, NS-nonsignificant; (D) Autosome vs. X-chromosomal transcripts m6A enrichment in different somatic cell types (left), and tissues (right). Mann-Whitney U test, **P value < 0.01, ****P value < 0.0001, NS-nonsignificant; MEF-mouse embryonic fibroblasts, NPC-neural progenitor cells, MSC-mesenchymal stem cells, maDC-mature dendritic cells, imDC-immature dendritic cells, regDC-regulatory dendritic cells.

### The extent of X to autosome dosage compensation differs between m6A methylated vs. non-methylated X-linked transcripts

Next, we were curious to know if m6A RNA methylation contributes to the maintenance of X to autosome dosage compensation as m6A were differentially enriched between X-linked vs. autosomal transcripts (Fig. 3). It is believed that less enrichment of m6A on X-linked transcripts increase their stability and thereby partly contribute to the maintenance of the X to autosome dosage equilibrium. To explore this further, we compared the X to autosome dosage compensation pattern between m6A methylated vs. non-methylated X-linked transcripts. To profile the X to autosome dosage compensation pattern, we calculated allelic X to autosome expression ratio in XEN, TSC and EpiSC. If there is X to autosome dosage compensation the active-X^Mol^:A^Mol^ ratio is expected to be > 1 and close to 2. Indeed, we find that active-X^Mol^:A^Mol^ ratio is ∼2 in XEN, and TSC cells, however, for EpiSC, we observed the ratio is slightly higher than 2 (Fig. 4A). We conclude that XEN, TSC and EpiSC undergo robust X to autosome dosage compensation in these cell lineages. To test further, we compared the overall allelic expression of autosomal and X-linked genes. We observed that the expression of X-linked genes from the active-X was significantly higher compared to the autosomal allelic expression in XEN, TSC and EpiSC, corroborating the upregulation of X-linked gene expression (Fig. 4B). Next, we asked if the extent of X to autosome dosage compensation differs between m6A methylated vs. non-methylated transcripts in XEN, TSC and EpiSC. To explore this, we profiled active-X^Mol^:A^Mol^ ratio by segregating the X-linked transcripts to m6A methylated vs. non-methylated. We find that active-X^Mol^:A^Mol^ ratio for X-linked transcripts without m6A methylation was significantly higher compared to the m6A modified X-linked transcripts, suggesting that the extent of X to autosome dosage compensation differs between m6A methylated vs. non-methylated X-linked transcripts (Fig. 4C). To test further, we profiled the allelic expression of autosomal transcripts and m6A methylated/non-methylated X-linked transcripts (Fig. 4D). We find that active-X expression of both m6A and non-m6A X-linked transcripts are higher compared to the allelic autosomal expression, indicating both classes of RNAs are upregulated (Fig. 4D). Interestingly, the expression of non-m6A X-linked transcripts tend to be higher compared to the m6A modified X-linked transcripts, corroborating the fact that the extent of upregulation differs between m6A vs. non-m6A X-linked transcripts (Fig. 4D). Taken together, we conclude that the degree of X to autosome dosage compensation correlates with the m6A methylation pattern of X-linked transcripts.

**Fig. 4:**
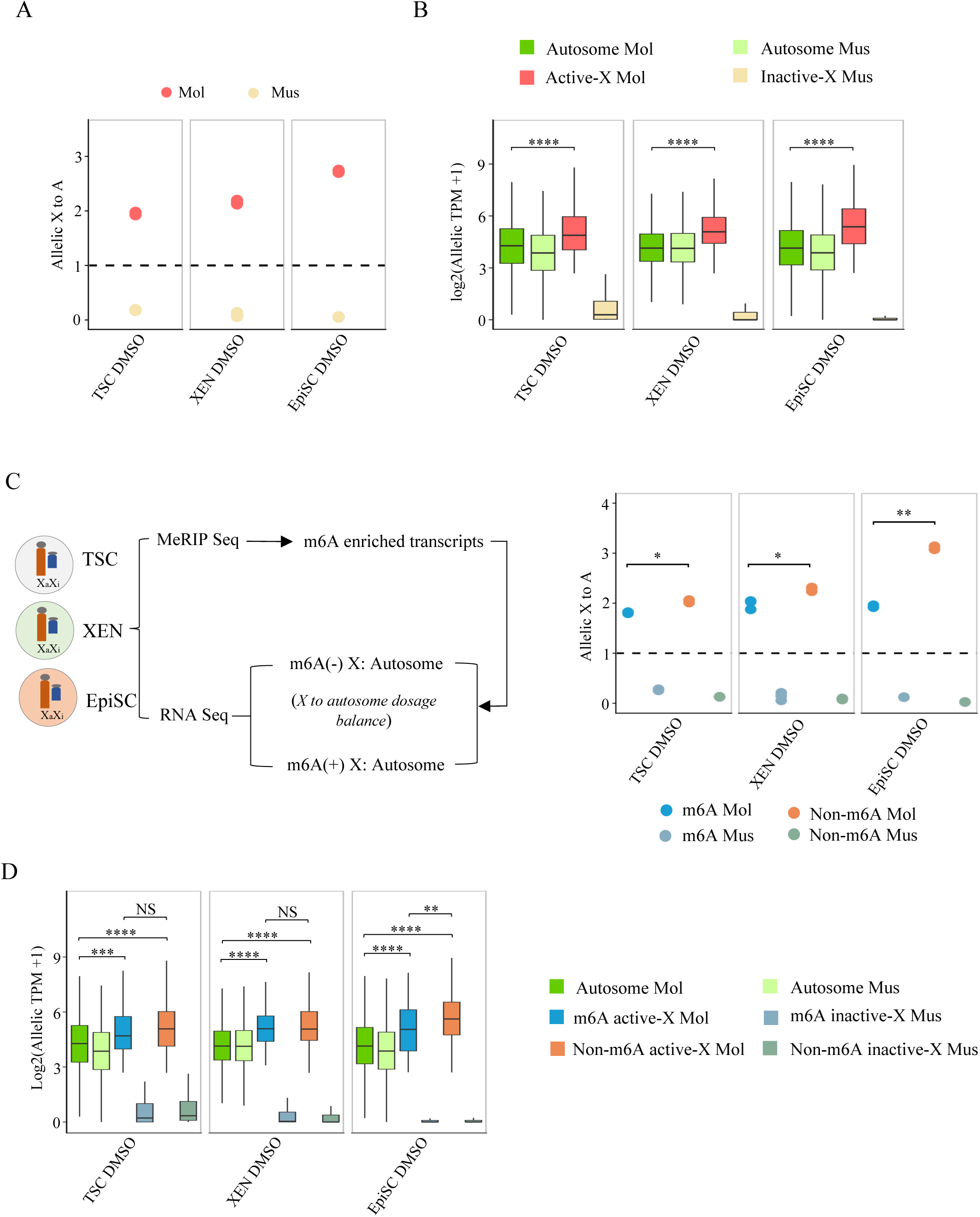
Profiling of X to autosome dosage compensation pattern of m6A and non-m6A methylated X-linked transcripts in TSC, XEN and EpiSC. (A) Plots showing allelic X:A ratio in DMSO treated TSC, XEN and EpiSC. (B) Boxplot showing the allelic expression of autosomal and X-linked genes in DMSO treated TSC, XEN, and EpiSC; Mann-Whitney U test; ****P value < 0.0001 (C) Schematic showing analysis workflow for profilling X:A ratio for m6A and non-m6A methylated X-linked transcripts (left); Allelic X:A ratio of m6A and non-m6A methylated X-linked transcripts in TSC, XEN and EpiSC (right); Two sample t-test; *P value < 0.05, **P value < 0.01. (D) Boxplot showing allelic expression of autosomal and m6A enriched or non-m6A enriched X-linked genes in DMSO treated TSC, XEN, and EpiSC; Mann-Whitney U test; **P value < 0.01, ***P value < 0.001, ****P value < 0.0001, NS – nonsignificant. TSC (n=2), XEN (n=3), and EpiSC (n=2).

### Depletion of m6A RNA methylation has minor effects in maintaining X to autosome dosage compensation

It is believed that while less enrichment of m6A on X-linked transcripts increases their stability, higher enrichment on autosomal transcripts reduces the stability of autosomal RNAs and thereby contributes to the maintenance of the X to autosome dosage balance. Therefore, we investigated if m6A RNA methylation contributes to the maintenance of the X to autosome dosage compensation in XEN, TSC and EpiSC. To explore this, we compared the allelic active-X^Mol^:A^Mol^ ratio between DMSO and inhibitor-treated (m6A-depleted) XEN, TSC and EpiSC cells (Fig. 5A). Interestingly, we found that depletion of m6A RNA methylation in XEN, TSC and EpiSC resulted in a slight downward shift in the active-X^Mol^:A^Mol^ ratio compared to DMSO control, however, the difference was not statistically significant in TSC and XEN cells (Fig. 5B). To test further, we compared the overall expression of autosomal and X-linked genes between DMSO vs. m6A-depleted XEN, TSC and EpiSC at an allelic resolution. We observed that expression of X-linked genes from the active-X was significantly higher compared to the autosomal allelic expression in inhibitor-treated cells despite the depletion of m6A RNA methylation (Fig 5C). Next, we tested if there were overall changes in allele-wise expression of X-linked and autosomal genes between DMSO vs. inhibitor-treated cells. Surprisingly, we did not observe any significant changes in expression of X-linked genes on the active-X chromosome between DMSO vs. inhibitor treated cells (Fig 5C). However, we found significant changes in autosomal allelic expression between DMSO and inhibitor-treated cells (Fig 5C). To explore further, we also compared active-X^Mol^:A^Mol^ ratio for m6A methylated and non-methylated X-linked transcripts in the DMSO vs. inhibitor-treated cells. Surprisingly, we did not observe any significant differences in active-X^Mol^:A^Mol^ ratio between DMSO vs. inhibitor-treated cells for m6A modified X-linked transcript (Fig. 5D). However, we noticed a downward shift for active-X^Mol^:A^Mol^ ratio for m6A non-methylated X-linked transcripts in inhibitor treated cells (Fig. 5D). However, the difference of active-X^Mol^:A^Mol^ ratio for m6A non-methylated X-linked transcripts between DMSO vs. inhibitor-treated cells was not significant in TSC and XEN (Fig. 5D). But, in case of EpiSC, we find that the difference is statistically significant (Fig. 5D). Additionally, we observed that despite of m6A depletion, both m6A methylated and non-methylated X-linked transcripts maintained higher allelic expression from the active-X compared to the autosomes, suggesting no effect on X to autosome dosage compensation (Fig. 5E). Taken together, we conclude that depletion of m6A RNA methylation has a minor impact on X-to-autosome dosage equilibrium in XEN, TSC and EpiSC.

**Fig. 5:**
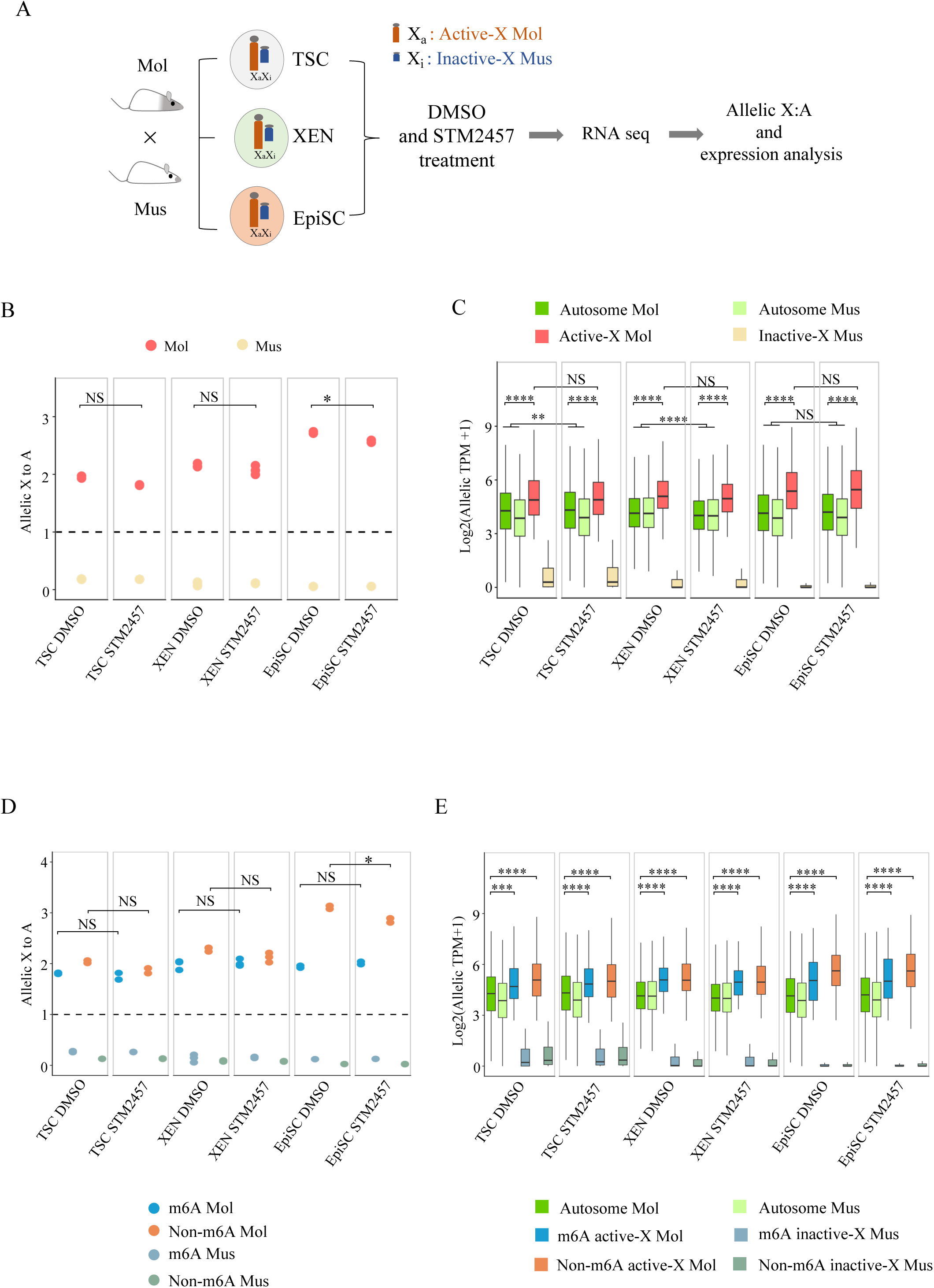
Effect of depletion of RNA m6A levels in the maintenance of X to autosome dosage in TSC, XEN and EpiSC. (A) Schematic outline showing allelic X to autosome ratio analysis in the DMSO and STM2457 treated TSC, XEN, and EpiSC. (B) Plots showing allelic X:A ratio in the DMSO vs. STM2457 treated TSC, XEN and EpiSC; Two sample t-test; *P value < 0.05, NS-nonsignificant; (C) Boxplot showing allelic expression of autosomal and X-linked genes in the DMSO and STM2457 treated TSC, XEN and EpiSC; Mann-Whitney U test; **P value < 0.01, ****P value < 0.0001; NS – nonsignificant. (D) Plots showing the allelic X:A ratio of m6A and non-m6A enriched X-linked transcripts in DMSO vs. STM2457 treated TSC, XEN and EpiSC; Two sample t-test; *P value < 0.05; NS – nonsignificant; (E) Boxplot showing allelic expression level of autosomal and X-linked genes (m6A and non-m6A enriched) in DMSO and STM2457 treated TSC, XEN and EpiSC; Mann-Whitney U test; ***P value < 0.001, ****P value < 0.0001. TSC (n=2), XEN (n=3), and EpiSC (n=2).

Next, we extended our investigation to test whether m6A depletion perturbs the X to autosome dosage equilibrium in other cell types. We compared the X:A ratio in wild-type (WT) vs. m6A-depleted Mettl3 knockout (KO) or knockdown (KD) ESCs, maDCs, MSCs, 3T3-L1 MEFs, brain and mouse liver cells through the analysis of available RNA-seq datasets (Geula et al., 2015; C.-X. Wang et al., 2018; H. Wang et al., 2019; S. Wang et al., 2023; Wu et al., 2018; Xu et al., 2021; Zhao et al., 2014). In most of the cases, we observed a slightly downward shift in the X:A ratio in Mettl3 knockout (KO) or knockdown (KD) cells compared to the WT cells. However, the differences were not significant in most of the cell types except ESC-E14Tg-2a and liver (Fig. 6A). Association of METTL14 is believed to be essential for the METTL3 to deposit m6A modification in RNA. Therefore, we analysed the impact of m6A depletion in Mettl14 knockout on the maintenance of an equilibrium X to autosome dosage from available ESC and NPC cell line datasets (Duda et al., 2021; Y. Wang et al., 2018b). Here, we observed a similar scenario where m6A depletion in Mettl14 KO cells only showed a downward trend in the X:A ratio compared to WT cells (Fig. 6B). However, we observed that the difference was significant in ESC (Fig. 6B). Taken together, our analysis shows that ablation or KD of m6A-writer enzymes only have minor effects in maintaining X to autosome dosage.

**Fig. 6:**
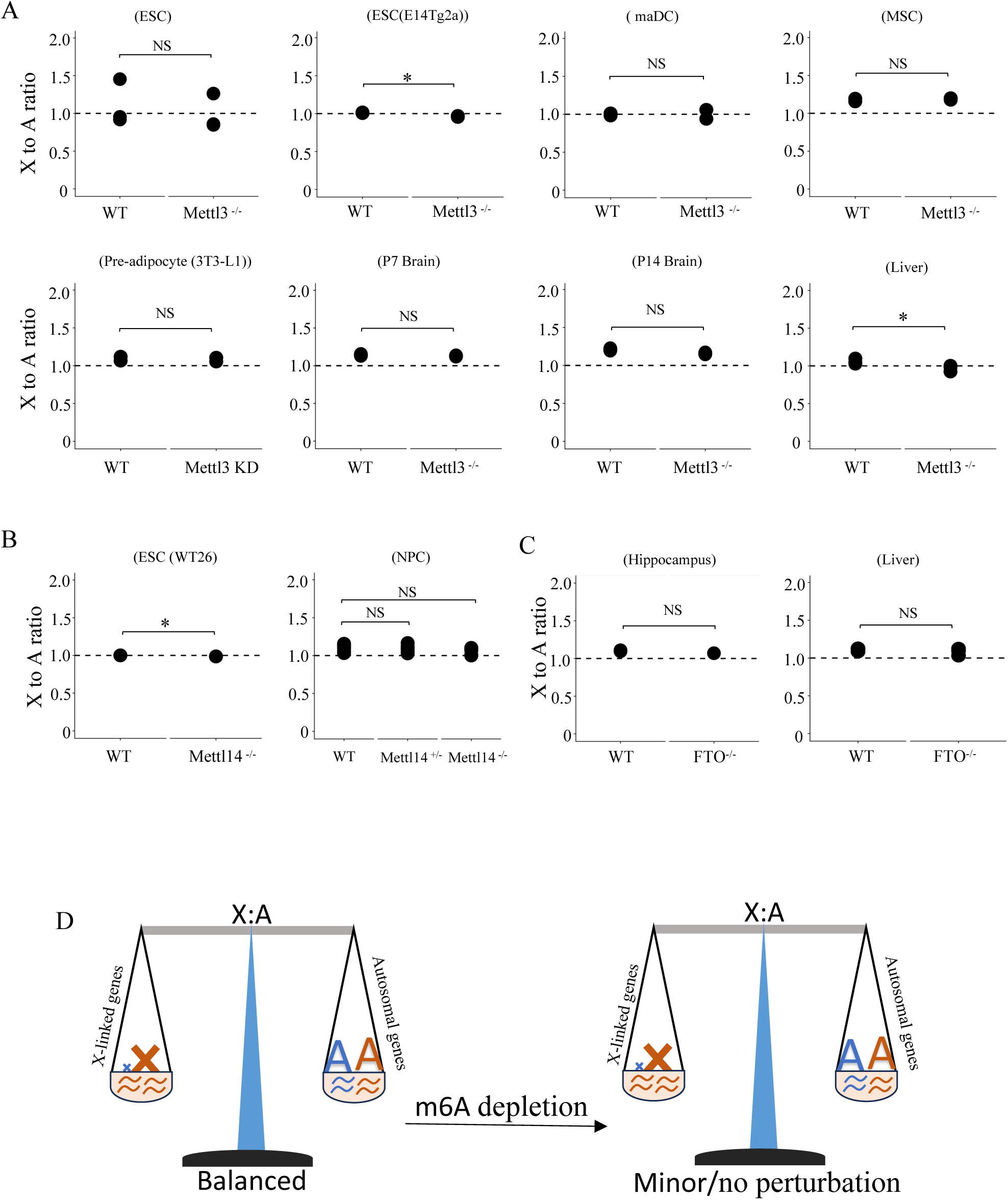
Effect of depletion of RNA m6A levels on X to autosome dosage balance in various cell types and tissues. (A) Comparison of X:A ratio between wild type (WT) and Mettl3 KO or KD cells and tissues; Two sample t-test; *P value < 0.05, NS – nonsignificant. (B) Comparison of X:A ratio between WT and Mettl14 KO cells; Two sample t-test; *P value < 0.05, NS – nonsignificant. (C) Comparison of X:A ratio between WT and FTO KO tissuesin hippocampus and liver; Two sample t-test; NS – nonsignificant. (D) Model showing global depletion of m6A level in cells can have a minor effect on X to autosome dosage compensation.

To get a better insight into the essentiality of m6A in the maintenance of optimum X:A dosage, we analysed the change in X:A ratio upon KO of m6A-eraser enzyme FTO. With perturbed m6A-erasers, autosomal transcripts would be more destabilized compared to X-linked ones, which can affect the overall X to autosomal dosage compensation. Through our analysis, we observed a more-or-less similar X:A ratio in both WT vs. FTO KO cells of the liver and hippocampus (Fig. 6C) (L. Li et al., 2017; Peng et al., 2019). Cumulatively, our analysis suggests that lower m6A enrichment in X-linked transcripts may only have a minor role to play in X to autosome dosage compensation.

## Discussion

In this study, we have dissected the role of m6A RNA methylation in dosage compensation in early embryonic lineages as well as somatic cells. First, we explored the role of m6A RNA methylation in the maintenance of X-linked gene silencing on the inactive-X in embryonic lineages with imprinted XCI (XEN, TSC) and random XCI (EpiSC). Surprisingly, we observed that METTL3 inhibition-mediated depletion of m6a RNA methylation in these cells does not perturb the inactive-X gene silencing, suggesting m6A RNA methylation is dispensable for the maintenance of X-linked gene silencing (Figs. 2). Our results are consistent with previous studies which showed that Mettl3 knockout or even segmental deletion of certain m6A-containing sites on *Xist* lncRNA in mESCs barely abrogates the gene-silencing efficiency during the initiation of XCI (Coker et al., 2020; Nesterova et al., 2019). However, there have been contrasting results in mESCs that show that *XIST* RNA methylation promotes X-linked gene silencing through facilitated binding of several protein complexes involved in gene repression(Patil et al., 2016). Also, work from another group have demonstrated that m6A sites on *XIST* RNA plays a pivotal role in silencing of X-linked genes on the inactive-X in HEK293T cells (Chang et al., 2022). However, a major limitation in both of their study was that they experimented on only a few X-linked genes. Taken together, our study demonstrates that m6A RNA methylation is dispensable for the maintenance of X-linked gene silencing on the inactive-X in XEN, EpiSC and TSC cells. However, one of the limitations of our study is that we performed inhibitor treatment for 12hrs to deplete the m6A level; therefore, in the future, prolonged depletion of m6A may provide better clarity into the requirement of m6A RNA methylation in the maintenance of X-linked gene silencing on the inactive X-chromosome. Nevertheless, our study provides insight on the role of m6A RNA methylation upon acute depletion which tends to eliminate the secondary effects that may arise due to deletions or knocking out the genes.

On the other hand, we demonstrate that during early embryogenesis, starting from the 2-cell stage to the blastocyst stage, X-linked transcripts are less m6A methylated compared to the autosomal transcripts (Fig. 3A). The trend of reduced m6A levels on X-chromosomal transcripts was consistent in different embryonic lineages such as XEN, EpiSC, TSC and ESC (Fig. 3B and 3C). Most of the other cell types we analysed such as MEFs, NPCs, MSCs, dendritic cells and liver tissues etc. also exhibited similar trends and other cell types (Fig. 3D). Taken together, we conclude that reduced enrichment of m6A modification on X-linked transcripts compared to the autosomal transcripts is consistent across most of the embryonic or somatic cell types that we have analysed. Previously, it was demonstrated that RNA-mediated regulation of differential transcript stability of X-chromosome vs. autosomes contributes to modulating the transcriptional dosage between monoallelic X-linked genes and biallelic autosomal genes(Deng et al., 2013). Indeed, a recent study has shown that preferential destabilisation of autosomal transcripts over X-linked transcripts is the result of higher m6A enrichment on autosomal transcripts in mESCs and different mouse and human tissue types. Moreover, they showed that the X-linked transcripts have evolved to bear lesser m6A motifs and thereby, have lesser m6A enrichment intrinsically compared to autosomal RNAs (Rücklé et al., 2023). Overall, our data support their findings and demonstrate the universality of the observation in different cellular contexts. Interestingly, we found that the extent or degree of X to autosome dosage compensation differs between m6A methylated and non-methylated X-linked transcripts in XEN, TSC and EpiSC (Fig. 4).

Next, we explored if m6A is required for the maintenance of X to autosome dosage compensation in XEN, TSC and EpiSC. Our analysis revealed that depletion of global m6A in XEN, TSC and EpiSC leads to a slight decrease in active-X^Mol^:A^Mol^ ratio, suggesting that m6A RNA methylation contribute partly for maintaining the X to autosome dosage in XEN, TSC and EpiSC (Fig. 5B). Indeed, a recent study has shown that the loss of m6A in ESC and other cell types leads to the reduction in X:A ratio (Ruckle et al. 2023). However, we find that the magnitude of the reduction in X:A ratio upon m6A depletion in XEN, TSC and EpiSC varied. While m6A depleted XEN and TSC showed a slight decrease in X:A ratio, EpiSC showed a significant reduction, suggesting m6A loss impact the X-to autosome dosage compensation in a lineage-specific manner (Fig. 5B). On the other hand, we found that there were no significant changes in X-linked gene expression between inhibitor and DMSO-treated cells, however, there were alterations in autosomal gene expression between the inhibitor and DMSO-treated cells, suggesting that the loss of m6A destabilizes the X to autosome dosage balance mainly through affecting the autosomal gene expression (Fig. 5C). Separately, although we observed significant global depletion of m6A upon inhibitor treatment for 12hr, we believe a more stringent strategy of prolonged m6A or stable m6A depletion may provide better clarity on the contribution of m6A RNA methylation in the maintenance of X:A dosage compensation in XEN, TSC and EpiSCs. Next, we extended our analysis to different embryonic and somatic cell types. Our analysis revealed that despite the depletion of m6A RNA methylation (∼70-99%) in Mettl3 and Mettl14 (m6A-writer) deletion or knockdown cells, there was no significant alteration in the X:A ratio in most of the cases (Fig. 6A and B). However, we would like to mention that in many cases, we observed a slight downward shift in the X:A ratio in m6a-depleted cells. Taken together, we demonstrate that m6A RNA methylation is required but has a minor role in the maintenance of X to autosome dosage compensation. On the other hand, gain of m6A enrichment in FTO (m6A-eraser) knockout hippocampus or liver cells did not alter the X:A ratio (Fig. 6C). It is worth mentioning that in some of the datasets that we have analysed, we do not have clarity about the efficiency of loss or gain of m6A enrichment such as Mettl3 knockdown in 3T3-L1, MEF cells (Zhao et al., 2014), Mettl14 knockout in WT26 ES cell lines (Duda et al., 2021) and FTO knockout in mouse liver (Peng et al., 2019). In summary, our study provides significant insight into the role of m6A RNA methylation in regulating dosage compensation. We demonstrate that m6a-RNA modification is dispensable for the maintenance of XCI, and has a minor contribution towards balancing the X to autosome dosage.

## Materials and Methods

### Data availability

RNA-seq and Me-RIP seq datasets generated in this study will be deposited to Gene expression omnibus (GEO) and the reference will be provided upon publication. Details about the previously available RNA-seq and Me-RIP seq dataset have been provided in Supplementary file 1.

### Cell culture

All cell lines (XEN, TSC and EpiSC) were grown and maintained at 37°C incubator in a humid atmosphere with 5% CO2. For growing XEN cells, culture dishes were coated with 0.2% gelatin (HiMedia, TCL059). Dulbecco’s modified eagle medium (DMEM) (HiMedia, AL007A) was used for culturing XEN cells with supplementation of 10% fetal bovine serum (FBS) (Gibco, 10270-106), Penstrep (Gibco, 15070063), L-Glutamine (Gibco, 25030081), MEM non-essential amino acids (NEAA) (Gibco, 11140-050) and β-Mercaptoethanol (Sigma, M3148).

EpiSCs were cultured on Matrigel (Corning, 354277) coated culture dish using Knock Out DMEM (Gibco. 10829018) supplemented with 20% knock out serum replacement (KSR) (Gibco. 10828028), 75U/ml Penstrep (Gibco, 15070063), 3 mM L-Glutamine (Gibco, 25030081), 1.5X MEM NEAA (Gibco, 11140-050), β-Mercaptoethanol (Sigma, M3148), 10ng/ml FGF2 (Prospect, CYT386) and 20ng/ml Activin A (SinoBiologicals, 10429 HNAH).

TSCs were cultured on gelatin (0.2%) coated culture dish using MEF (mouse embryonic fibroblast) conditioned TSC medium. TSC medium consisted of RPMI (PanBiotech, P04-16500), L-Glutamine, Penstrep (Gibco, 15070063), MEM NEAA (Gibco, 11140-050), β-Mercaptoethanol (Sigma, M3148) and sodium pyruvate. To the finally prepared TSC medium 37.5ng/µl FGF4 (Peprotech, 45057) was added prior to use. For the preparation of MEF conditioned media, MEF cells were cultured with TSC medium in a gelatin-coated culture dish.

### METTL3 inhibitor treatment

To inhibit METTL3 in XEN, EpiSC and TSC, cells were seeded a day before the treatment. Next day, cells were treated with 50 µM of STM2457 inhibitor (STORM THERAPEUTICs) using the respective culture media for 12 hours. STM2457 inhibitor solution was prepared by dissolving in DMSO (Sigma, A2438). An equivalent volume of DMSO alone was used as a vehicle and treated for 12 hours in parallel to the inhibitor treatment. Following the treatment, cells were harvested in TRIzol reagent (Life technologies 15596-026).

### Total RNA isolation

Total RNA from all the cell lines (XEN, TSC and EpiSC) were isolated using TRIzol reagent (Life technologies 15596-026) following manufacturer’s instructions and resuspended in ultrapure H2O (Invitrogen, 10977). The concentration and integrity of all the RNA samples were measured using Nanodrop (Thermofisher Scientific) and running through 1% agarose gel respectively.

### LC MS/MS

For the liquid chromatography-tandem mass spectrometry (LC-MS/MS) experiment, mRNA from XEN, TSC and EpiSC was used. First, to remove the genomic DNA contamination, total RNA was treated with Turbo DNase (Invitrogen, AM2238) at 37°C for 45 mins, followed by purification by standard phenol-chloroform isoamyl alcohol method and resuspended in ultrapure H_2_O. Next, mRNA was isolated from the DNase-treated total RNA using Dynabeads Direct mRNA purification kit (Invitrogen, 61011) following the manufacturer’s instruction. To prepare the sample for LC-MS/MS, mRNA was hydrolysed in a buffer containing 10U of nuclease P1 (NEB, M0660S), NaCl and ZnCl2 for 2 hours with agitation at 800 rpm for 30s every 5 min at 37°C in a thermomixer (Thermofisher Scientific). Next, NH4CO3 (Sigma, A6141) and alkaline phosphatase (NEB, M0525S) were added and incubated for an additional 2 hours. Tris-HCl was added to stop the reaction and centrifuged at 16000g for 30 mins at 4°C. The supernatant was collected and injected into C18 reverse phase column coupled to Shimadzu triple quadrupole (QQQ) mass spectrometer in positive electrospray ionization method. The nucleosides were identified based on the nucleoside to base ion transitions of 268-to-136 for A and 282-to-150 for m6A. Nucleosides were quantified from the calibration curve generated from pure nucleosides of A (Sigma, A9251) and m6A (Abcam, ab145715).

### RNA sequencing and analysis

For RNA-seq, total RNA was isolated using TRIzol method as mentioned above. RNA quantity was checked on Qubit fluorometer (Thermofisher, Q33238) using RNA HS assay kit (Thermofisher, Q32851) following the manufacturer’s instructions. The purity and integrity of RNA were checked using Tapestation 4150 (Agilent) using HS RNA screen tape, respectively. RNA samples with RIN > 8 were used for the library preparation using TruSeq stranded total RNA kit (Illumina, 15035748). Yield of libraries was quantified in Qubit 4.0 fluorometer (Thermofisher, Q33238) using DNA HS assay kit (Thermofisher, Q32851) and insert size was determined on Tapestation 4150 (Agilent) using D1000 screentapes (Agilent, 5067-5582). Libraries were finally sequenced on Illumina NovaSeq 6000 platform with paired-end (2 x150) chemistry. For analysis of RNA-seq data, the quality of reads were checked using FastQC (v0.12.1; https://www.bioinformatics.babraham.ac.uk/projects/fastqc/). Next, adapters were removed using Trim_galore (v0.6.10); (http://www.bioinformatics.babraham.ac.uk/projects/trim_galore/). Ribosomal reads were removed by ribodetector (v0.2.7) (Z. L. Deng et al., 2022). RNA-seq reads were mapped to mouse reference genome GRCm39 using STAR (v2.7.10a) aligner (Dobin et al., 2013). Gene wise featureCounts (v2.0.1) was used to generate counts and normalized to TPM (Liao et al., 2014).

### Allele-specific RNA-seq analysis

Allelic expression profile of X-linked and autosomal genes was obtained through analysis of the RNA-seq data using an allele-specific expression analysis pipeline based on strain-specific SNPs as described previously (Ayyamperumal et al., 2024; Naik et al., 2022, 2024). We obtained strain-specific SNPs (Mus musculus musculus (Mus) and Mus musculus molossinus (Mol)) from the Mouse genome project (https://www.sanger.ac.uk/science/data/mouse-genomes-project). Next, we created strain-specific in silico reference genomes using the variant calling file (VCF) tool (Danecek et al., 2011). Basically, we generated strain-specific in silico reference genomes by incorporating strain-specific SNPs into the GRCm39 reference genome. Next, transcriptomic reads were mapped to the strain-specific in silico reference genomes using STAR (-- outFilterMultimapNmax 1). Following the mapping of the reads, SNP-wise counts were generated. After generating SNP-wise read counts, we applied the following filters to remove any false positives in our allelic counts. First, SNPs with minimum read counts of 10 per SNP site were only selected for further analysis. Next, to obtain the allelic count for individual genes, we considered only those genes that had at least two SNPs that qualified the above SNP threshold. Additionally, the allelic read counts for individual genes were deduced by taking an average of SNP-wise reads. Finally, the allelic ratio was calculated using the following formula: (Mol-allele or Mus-allele)/(Mol-allele+Mus-allele).

### X to A ratio analysis

X to autosome expression ratio was estimated following the pipeline as described previously (Naik et al., 2022; Pacini et al., 2021). First, we defined gene sets for both X-chromosome and autosomes by excluding lowly expressed genes (<5 TPM) from our analysis. Similarly, highly expressed genes were removed from autosomes and X chromosomes by applying a 95-percentile threshold. Using these filtered gene set, X:A ratio was calculated using the bootstrapping approach. Basically, X-linked gene expression (TPM) was divided with the same number of autosomal genes selected randomly and was repeated 1000 times to mitigate a large number of autosomal genes compared to the small number of X-chromosomal genes. Finally, X to A ratio was obtained from the median value of 1000 repeats.

### MeRIP - sequencing

For performing MeRIP in XEN, TSC and EpiSCs, EpiQuik CUT&RUN m6A RNA enrichment kit (EpiGenTek# P-9018) was used. In brief, total RNA was isolated as described above and MeRIP was performed according to the manufacturer’s protocol of the EpiQuik CUT&RUN m6A RNA enrichment kit with slight modifications. Total RNA was DNase-treated before the immunoprecipitation. Immunoprecipitation (IP) was performed using the m6A antibody (provided in the kit) coupled with affinity beads for overnight at 4°C. After IP, the RNA was purified and subjected to library preparation using the NEB Next Ultra II Directional RNA Library kit (NEB# E7765) and sequenced using Illumina NovaSeq 6000 using 2×150 paired-end chemistry.

### MeRIP-seq data analysis

First, the quality of MeRIP-seq reads were checked using FastQC (v0.12.1; https://www.bioinformatics.babraham.ac.uk/projects/fastqc/). Next, reads were trimmed for adapters and low-quality bases using Trim_galore (v0.6.10; (http://www.bioinformatics.babraham.ac.uk/projects/trim_galore/). Additionally, ribosomal reads were removed by ribodetector (v0.2.7) (Z. L. Deng et al., 2022). After all these quality checks, MeRIP-seq reads were mapped to the mouse reference genome GRCm39 using STAR (v2.7.10a) aligner (Dobin et al., 2013). Bigwig files were generated using deeptools bamCoverage (v3.5.5) with --normalizeUsing BPM and visualised by integrated genomic viewer (IGV). Gene wise featureCounts (v2.0.1) was used to get counts and normalized to TPM (Liao et al., 2014). Input genes which have >0.5 TPM was used for further analysis to identify m6A enrichment IP over Input and generated log2 (IP TPM/Input TPM +1) values. m6A peaks were identified using MACS2 (v 2.2.9.1)(Zhang et al., 2008) peak caller function “callpeak” with the following parameters: ‘--nomodel’, ‘--keep-dup all’, ‘-g 2654621783’ and statistical cutoff q 0.05 value. Significant peaks were intersected with the exonic region of genes using bedtools (v2.30.0) (Quinlan & Hall, 2010)

### Statistical analysis and plots

Plots were generated using R packages ggplot2, Complex Heatmap and python package pyGenomeTrackstracks (Lopez-Delisle et al., 2021). For statistical analysis, we used two-sample t-tests and Mann-Whitney U test.

## Author’s contribution

Conceptualization: SG, HCN, RB and AS. Supervision and funding: SG. Funding: Experiments, data analysis, methodology and resources: HCN, RB, MNM and KS. Writing first draft: AS, RB and HCN. Editing the manuscript: SG, AS, HCN and RB. The final manuscript was approved by all the authors.

## Acknowledgement

We thank Prof. Sundeep Kalantry, University of Michigan for providing cell lines used for our experiments in this study. This study is supported by the Department of Biotechnology (DBT), Govt. of India grant (BT/PR30399/BRB/10/1746/2018), DST-SERB (CRG/2019/003067), DBT-Ramalingaswamy fellowship (BT/RLF/Re542 entry/05/2016) and Infosys Young Investigator grant award to SG. RB acknowledge Council of Scientific and Industrial Research (CSIR), India for the fellowship. AS acknowledges MOE and UGC for the fellowship.

